# Injectable diblock copolypeptide hydrogel provides platform to maintain high local concentrations of taxol and local tumor control

**DOI:** 10.1101/688762

**Authors:** Matthew C. Garrett, Timothy M. O’Shea, Alexander L. Wollenberg, Alexander M. Bernstein, Derek Hung, Brittany Staarman, Horacio Soto, Timothy J. Deming, Michael V. Sofroniew, Harley I. Kornblum

**Affiliations:** Department of Neurosurgery, University of Cincinnati College of Medicine, Cincinnati OH 45267, USA; Department of Neurosurgery David Geffen School of Medicine, University of California Los Angeles, Los Angeles CA 90095-0119, USA; Department of Chemistry and Biochemistry, University of California Los Angeles, Los Angeles CA 90095-1569, USA; Department of Neurobiology, David Geffen School of Medicine, University of California Los Angeles, Los Angeles CA 90095-1763, USA; Department of Bioengineering, University of California Los Angeles, Los Angeles CA 90095-1600, USA; Departments of Psychiatry and Biobehavioral Sciences, Pharmacology, Pediatrics and the Semel Institute for Neuroscience and Human Behavior, David Geffen School of Medicine, University of California Los Angeles, Los Angeles CA 90095-1655, USA

## Abstract

**Introduction:** Surgical resection and systemic chemotherapy with temozolomide remain the mainstay for treatment of glioblastoma. However, many patients are not candidates for surgical resection given inaccessible tumor location or poor health status. Furthermore, despite being first line treatment, temozolomide has only limited efficacy.

**Methods:** The development of injectable hydrogel-based carrier systems allows for the delivery of a wide range of chemotherapeutics that can achieve high local concentrations, thus potentially avoiding systemic side effects and wide-spread neurotoxicity. To test this modality in a realistic environment, we developed a diblock copolypeptide hydrogel (DCH) capable of carrying and releasing paclitaxel, a compound that we found to be highly potent against primary gliomasphere cells.

**Results:** The DCH produced minimal tissue reactivity and was well tolerated in the immune-competent mouse brain. Paclitaxel-loaded hydrogel induced less tissue damage, cellular inflammation and reactive astrocytes than cremaphor-taxol (typical taxol-carrier) or hydrogel alone. In a deep subcortical xenograft model, of glioblastoma in immunodeficient mice, injection of paclitaxel-loaded hydrogel led to a high local concentration of paclitaxel and led to local tumor control and improved survival. However, the tumor cells were highly migratory and were able to eventually escape the area of treatment.

**Conclusions:** These findings suggest this technology may be ultimately applicable to patients with deep-seated inoperable tumors, but as currently formulated, complete tumor eradication would be highly unlikely. Future studies should focus on targeting the migratory potential of surviving cells.

## 1. INTRODUCTION

Glioblastoma accounts for 40% of primary brain tumors and results in over 15,000 deaths a year in America alone[1]. Despite maximal therapy including surgical resection, average survival is only 20 months [3]. Some patients are not candidates for surgery due to poor health, deep or bilateral location or proximity to eloquent structures [5, 6]. Without cytoreductive surgery, the average survival is four months[3].

In addition to cytoreductive surgery, current standard of care also includes temozolomide and radiation. Temozolomide treatment has a small but statistically significant improvement in survival[7]. However, it also induces mutations leading to more aggressive recurrent tumors[8]. Further, treatment with temozolomide can result in enhanced resistance by demethylatation of the MGMT promoter[9]. There are many promising chemotherapy options that show improved potency over temozolomide without the genotoxic damage. However, often these drugs are limited by their pharmacokinetics, inability to cross the blood-brain barrier or difficulties and side effects with systemic administration[10].

To address these limitations, drug releasing implants are being developed, most often composed of biodegradable polymers, that may provide improved local delivery of promising chemotherapeutics in the CNS [11]. While multiple studies have evaluated the ability of different carriers to deliver chemotherapeutic payloads to glioblastoma cells, the majority of these studies have tested them in sub-optimal conditions using either in vitro [12] or subcutaneous flank environments [13], or testing them on suboptimal, multi-passaged glioma cell lines that often do not recapitulate the migratory capacity of patient tumors [14] [15] [16]. Fully understanding the potential of these vehicles requires testing in realistic in vivo environments. To this end, we have constructed diblock copolypeptide hydrogels (DCH) capable of delivering paclitaxel to a patient-derived gliomasphere model of GBM in a mouse brain. We found that paclitaxel-loaded hydrogels led to local control and improved survival in this mouse model, but surviving cells were able to migrate out of the treatment zone.

## METHODS

### Gliomasphere culture and in vitro proliferation assay

Cancerous stem cells can be derived from brain tumors and propagated in vitro as neurosphere-like gliomaspheres [17] [18]. For this study, we utilized the sample HK308, a gliomasphere line derived from a recurrent GBM in a 50-year-old male as previously described [17, 18]. The culture and the original tumor sample have an amplification of the epidermal growth factor receptor gene, and express the EGFRvIII mutation. MGMT was unmethylated in the original tumor sample as reported by the neuropathologist [17] [18]. For cell proliferation experiments used to determine potency of chemotherapeutic agents, gliomaspheres were initially plated in a 6 well plate using 2ml of serum-free media (DMEM F12 +B27+bFGF+EGF+heparin) at a concentration of 100,000 cells/ml. Chemotherapeutic drugs were solubilized in either aqueous or DMSO-based solutions at various concentrations and applied to the proliferating gliomasphere cells for seven days. On the seventh day the total number of viable cells were counted using Trypan blue with the Growth Rate (GR) value for the various drug concentrations determined by estimating the exponential growth kinetics over the seven days and comparing these to an untreated control using methods described previously [17]. Gliomasphere-derived hGBM cells used for *in vivo* mouse studies were transduced with a 3^rd^ generation, self-inactivating lentiviral construct expressing GFP and firefly luciferase [19] by dissociating 100,000 hGBM cells in 2mL of serum free media and adding 1mL of lentivirus and incubating for 3 days. Infected cells were purified by cell sorting for GFP and expanded *in vitro*.

### Hydrogel Synthesis and Fabrication

The diblock copolypeptide, K_180_A_40_ was the DCH used in this study, and was synthesized using techniques described in detail previously [20]. Briefly, within an inert, dinitrogen filled glovebox environment a solution of N_ε_-carbobenzyloxy-L-lysine N-carboxyanhydride ((Z-L-lysine NCA) in THF solvent was polymerized upon addition of the transition metal initiator, Co(PMe_3_)_4_. The polymerization of the N_ε_-carbobenzyloxy-L-lysine block was allowed to proceed for approximately 45 minutes and was monitored for completeness by FTIR before the subsequent addition of a solution of L-alanine NCA in THF. The consumption of the second NCA was also allowed to proceed for 45 minutes before the reaction was removed from the glovebox for subsequent deprotection of the N_ε_-carbobenzyloxy-L-lysine residues by HBr in an acetic acid/TFA solution followed by polymer precipitation in diethyl ether. The precipitated polymer was re-dissolved in nonpyrogenic DI water and then dialyzed within a 2000 Da MWCO dialysis bag exhaustively against NaCl for two days and then nonpyrogenic DI water for two days before being lyophilized. Paclitaxel was loaded into the DCH by dissolving and mixing the DCH and paclitaxel in equal mass in an 80/20 Methanol/water solution and then drying under vacuum overnight to remove the solvent. The dry DCH/paclitaxel powder was resuspended in nonpyrogenic DI water at 3 wt% for subsequent *in vitro* and *in vivo* evaluation.

### In vitro evaluation of release of paclitaxel from DCH

For the *in vitro* paclitaxel release assay, 100 µL of 3 weight-percent (wt%) DCH loaded with paclitaxel (DCH-paclitaxel) was injected into pre-hydrated 20,000 MWCO Slide-A-Lyzer dialysis cassettes before being placed in a sink of 100 mL of 5% Fetal Bovine Serum (FBS) containing 1xPBS. A cassette contacting 100 µL of paclitaxel in DMSO was used as a control. The complete incubation media was collected and replaced on days 1, 3, 7, 14, 21, 28 and 42. To analyze the amount of released paclitaxel, 50 mL of the incubation media from each time point was extracted with four separate 50 mL ethyl acetate washes. The extracted organic layer was evaporated to dryness under vacuum, resolubilized in acetonitrile and filtered with a 0.2 µm syringe filter prior to analysis by HPLC. HPLC analysis of paclitaxel concentration was performed using methods adapted from a previous report [21]. Specifically, 10 µL of release assay samples of unknown paclitaxel concentration and paclitaxel standards were run on a C18 150 × 4.6 mm I.D. Agilent column using an isocratic 50/50 Acetonitrile/Water mobile phase and detected by UV absorption detection at a wavelength of 227 nm.

### Surgery and Hydrogel implantation

Biocompatibility evaluations of DCH and Cremphor vehicle with and without paclitaxel were performed using adult male and female C57Bl6 mice aged 8-14 weeks. For gliomasphere-derived hGBM cell injections, Non-obese Diabetic/ Severe Combined Immunodeficiency/Gamma Null (NSG) mice were used. All surgical procedures performed within this paper were evaluated by the UCLA animal research committee (ARC) and described within an approved ARC protocol. Mice were anesthetized using isoflurane and a craniotomy was performed by drilling a rectangular exposure window in the bone with a high-speed dental drill. For biocompatibility evaluations, DCH or vehicle were injected stereotactically into the center of the caudate putamen nucleus using the target coordinates of 0.5 mm anterior to Bregma, 2 mm lateral to Bregma and a depth of 3.0 mm below the cortical surface. A 2 μL volume of DCH was injected using a pulled glass micropipettes ground to a beveled tip with 150–250 μm inner diameter attached via specialized connectors and high-pressure tubing to a 10 μL syringe that was mounted to a stereotaxic frame and controlled by an automated microdrive pump. For studies on hGBM cells, a 2µL suspension of 100,000 hGBM cells was injected first into the striatum using the target coordinates of 0.5 mm anterior to Bregma, 2 mm lateral to Bregma and a depth of 3.0 mm below the cortical surface. This first injection was followed immediately by a second injection of 2 µL of DCH placed directly above using the target coordinates of 0.5 mm anterior to Bregma, 2 mm lateral to Bregma and a depth of 2.0 mm below the cortical surface. Postoperative care and analgesia was administered preoperatively as well as for two days following surgery to alleviate pain.

### IVIS Spectrum *in vivo* Imaging

The gliomasphere cell proliferation in vivo within the brain was monitored at discrete time intervals using non-invasive bioluminescence imaging on an IVIS® Spectrum in vivo imaging system. Mice received intraperitoneal injections of 200 μL of luciferin and were scanned 5 minutes later with a 10 second exposure. Results were listed as flux (photons/second) through the area of interest.

### Animal Perfusions and Histology

At specific predetermined times after hydrogel/cell injection, or when tumors reached a terminal size requiring euthanasia, animals were given a lethal dose of pentobarbital and a transcardial perfusion was performed with heparinized saline and 4% paraformaldehyde. Brains were excised, post-fixed in 4% paraformaldehyde for 5 hours and then stored in 30% sucrose. Brains were sectioned along the coronal plane using a cryostat. Tissue sections (40 μM) were stained using standard fluorescent immunohistochemical techniques as previously described [22] to visualize host tissue, injected gliomaspheres cells and hydrogel deposit localization. Primary antibodies were: rabbit anti-GFAP (1:1000; Dako, Carpinteria, CA); rat anti-GFAP (1:1000, Zymed Laboratories); rabbit anti-Iba1 (1:1000, Wako Chemicals, Richmond VA); rat anti-CD68 (1:100; AbD Serotec, Biorad, CA). Fluorescence secondary antibodies were conjugated to: Alexa 488 (green) or Alexa 405 (blue) (Molecular Probes), or Cy3 (550, red) or Cy5 (649, far red) all from (Jackson Immunoresearch Laboratories). Nuclear stain: 4’,6’-diamidino-2-phenylindole dihydrochloride (DAPI; 2ng/ml; Molecular Probes). Sections were cover-slipped using ProLong Gold anti-fade reagent (InVitrogen, Grand Island, NY). Sections were examined and photographed using deconvolution fluorescence microscopy and scanning confocal laser microscopy (Zeiss, Oberkochen, Germany).

## RESULTS

### Paclitaxel is more potent than Temozolomide

Temozolomide is available in an oral formulation, is well tolerated, and is able to cross the blood-brain barrier making it a convenient therapeutic option. However, temozolomide has only modest efficacy in vitro[23] and in vivo [7]. Given space limitations the ideal candidate chemotherapeutic agent would have both high efficacy and high potency. We tested both temozolomide and paclitaxel at various doses against our patient-derived gliomasphere line (HK308). Both of the two candidate drugs were effective at suppressing the proliferation of the gliomaspheres in a concentration dependent manner over the seven-day evaluation period **(Figure 1A)**. By applying growth rate (GR) inhibition calculations[24], we quantified the relative effectiveness of each drug’s capacity to suppress cell growth in the endpoint assays. The GR_50_ value (representing the drug concentration at which the cell proliferation rate is reduced by half), was less than 1 nM (the lowest dose tested) for paclitaxel, and was approximately 0.5 mM for temozolomide. Based on these results, paclitaxel was at least 500,000 times more potent on our human patient derived gliomasphere cell line than temozolomide and was considered to be the more appropriate candidate for further exploration in subsequent *in vivo* studies.

**Figure 1.**
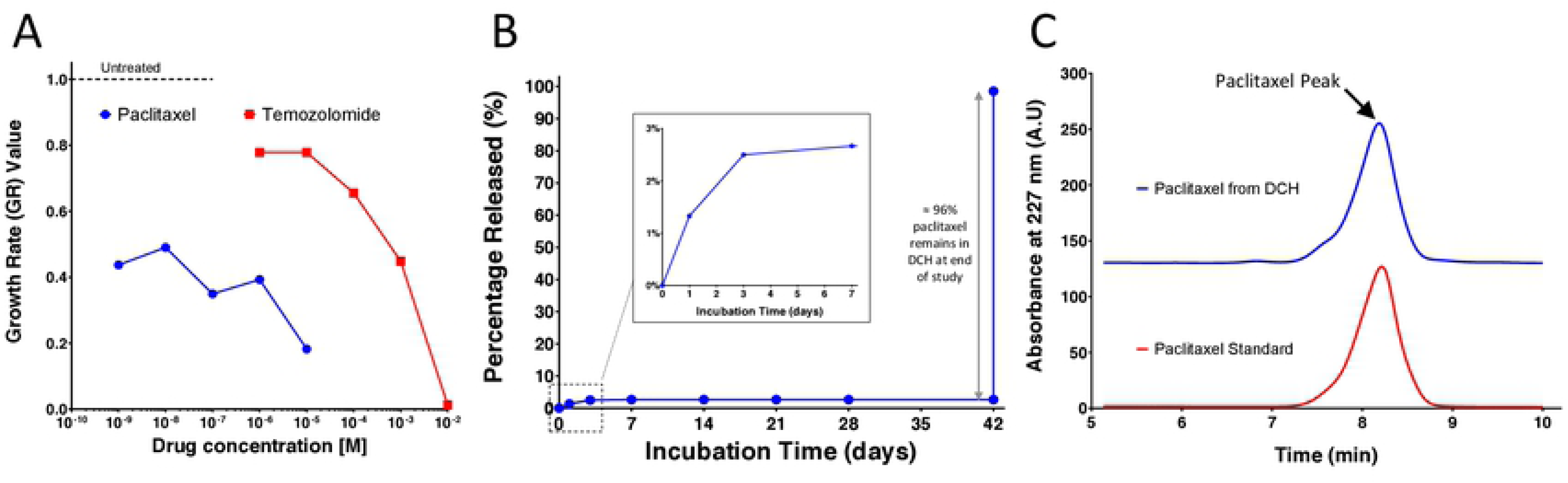
in vitro evaluation of DCH-paclitaxel system. **A.** On a per weight basis, paclitaxel is 500,000 times more potent than temozolomide at suppressing the growth of human gliomaspheres over a 7-day period *in vitro*. Growth rate (GR) inhibition calculations^27^ were used to quantify the relative effectiveness of various concentrations of each drug and normalized to untreated controls. **B.** Cumulative release curve for paclitaxel loaded into the DCH over a 6 week incubation in vitro. A 1000 fold sink of 5% FBS in 1XPBS was used. An initial burst release of paclitaxel of over the first three days was noted and while 96% of drug was recovered at the end of the 6 week incubation period. **C.** HPLC chromatograms show that paclitaxel recovered from the DCH at the end of the 6 week in vitro incubation was chemically identical to freshly prepared drug.

### DCH-Paclitaxel maintains a high local concentration over a prolonged period

The ideal chemotherapeautic agent should maintain high local concentrations and not become diluted through diffusion. To determine *in vitro* drug release kinetics we prepared a 3 wt% K_180_A_40_ DCH at a drug concentration of 3% w/v and placed the gel in a dialysis cassette suspended in an incubation buffer with 5% v/v fetal bovine serum (FBS) to simulate the brain environment. The incubation media was replaced at discrete time points and the concentration of paclitaxel was evaluated by HPLC following organic extraction. A solution of paclitaxel in DMSO at the same concentration was also incubated within a separate dialysis cassette over the same time course to serve as a control. The paclitaxel stayed inside the DCH matrix throughout the six week period indicating that the paclitaxel does not diffuse out of the matrix but is rather only released as the gel is degraded (Figure 1B). After the experiment, nearly all of the paclitaxel was recovered and found to have retained its chemical signature and biological activity (Figure 1C).

### DCH-Paclitaxel maintains a high local concentration over a prolonged period

To test the CNS biocompatibility of the DCH vehicle, we injected the K_180_A_40_ DCH with and without paclitaxel into the caudate putamen of healthy adult mice in the absence of hGBM cells and evaluated the general foreign body response using standard immunohistochemistry markers. For comparison, we injected an equivalent volume of 50% v/v Cremophor EL (polyoxyethylated castor oil) in ethanol with and without paclitaxel in separate cohorts of mice. Cremophor EL is used as the surfactant vehicle for paclitaxel within the standard Taxol formulation approved clinically for intravenous administration of the drug. Animals tolerated all four formulations well after injection and there were no observable adverse health events throughout the duration of the study. At one week post injection, which was previously characterized as the time of maximum foreign body response to DCH alone (without cargo molecules) [25], brain tissue was evaluated at and around the injection site using immunohistochemical markers for macrophages and activated microglia (CD68, IBA-1) and astrocyte reactivity (GFAP) (**Figures 2A-D**). To quantify the extent of the foreign body response, we characterized the intensity of immunohistochemical staining across a radial distance of 1 mm originating from the center of the injection (**Figure 2E,F**). To establish a cumulative measure of immunostaining across this circular tissue injection zone, we calculated the area under the curve (AUC) on the intensity plots (**Figure 2F**).

**Figure 2.**
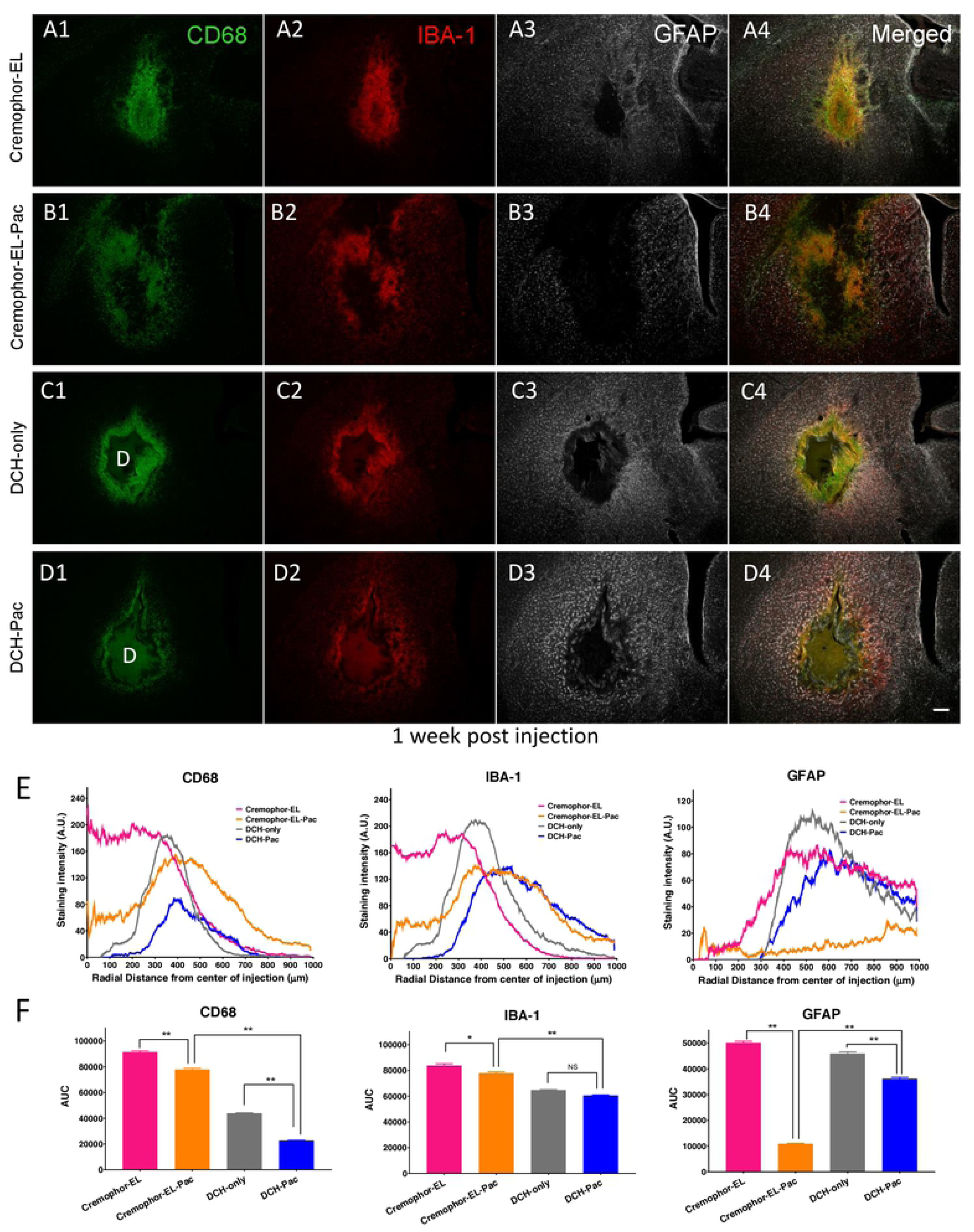
DCH-paclitaxel exhibits good biocompatibility in healthy CNS in contrast to paclitaxel in Cremophor EL vehicle. **A-D.** Images of caudate putamen at 1 week after injection into healthy, uninjured tissue of Cremophor EL vehicle (**A**), Cremophor EL + paclitaxel (**B**), DCH only (**C**), or DCH + paclitaxel (**D**), showing single channel and merged multichannel immunofluorescence for multiple markers of inflammation and gliosis, CD 68, IBA-1 and GFAP. Scale bar, 200 µm for all images, D = DCH depot. **E.** Quantification of immunofluorescence intensity for each treatment group across a radial area of 1 mm originating from the center of the injection (n=3 per stain per treatment). **F.** Area Under the Curve (AUC) calculations for the various immunofluorescence intensity traces from E. provide a single measure of cumulative staining within the 1 mm radial field. The DCH-paclitaxel system showed markedly and significantly less staining intensity, indicative of a more favorable foreign body response, compared with paclitaxel administered using the Cremophor EL vehicle * *p* < 0.01 or ** p < 0.001 (ANOVA with post-hoc Tukey’s multiple comparisons test).

The Cremophor EL vehicle treated group exhibited a small, localized core of densely packed CD68- and IBA-1-positive cells at the injection center, surrounded by a border of compact astrocyte reactivity typical of that around focal CNS lesions (**Figures 2A**). Compared with other treatment groups, the Cremophor EL vehicle showed the largest total of CD68-positive, IBA-1-positive and GFAP-positive cells within the field of analysis. By comparison, the inclusion of paclitaxel within this Cremophor EL vehicle resulted in a larger and more diffuse volume of tissue disruption and inflammation but significantly fewer total CD 68-positive and IBA-1-positive cells within the analysis field (**Figures 2B,F**). Interestingly, in the Cremophor EL plus paclitaxel group there was the appearance of increased microglia reactivity within the preserved tissue outside the direct lesion/disrupted tissue field suggesting soluble paclitaxel had reached these tissue regions and was mildly stimulating this cell population and/or that these cells were responding to cues from cells receiving paclitaxel in the core of the injection (**Figure 2B**). Notably, the injection region in the Cremophor EL plus paclitaxel group exhibited a remarkable and statistically significant absence of GFAP-positive cells, such that there was no compact border of reactive astrocytes in this group (**Figure 2B,F**).

Compared to the Cremophor EL treated groups, the DCH injected groups showed a milder and more focal foreign body response (**Figures 2C-F**), and in particular, there was significantly less CD68 immunoreactivity in the DCH and DCH plus paclitaxel groups (**Figures 2E,F**). The DCH-paclitaxel group also exhibited significantly less CD68 staining around the DCH boundary compared to the DCH only control (**Figures 2C-F**). Host neuronal viability was well preserved around the deposit margins, with only a limited radial zone of about 150 µm radially becoming depleted of NeuN positive neurons after 4 weeks of paclitaxel exposure (**Figure 3B**). By contrast paclitaxel delivered using Cremophor-El caused more profuse NeuN positive neuron loss around the injection site (**Figure 3C**) as well as a considerable depletion of GFAP positive astrocytes (**Figure 2B**). These findings provide strong evidence that the DCH-paclitaxel system caused less damage to host tissue compared to the standard paclitaxel in Cremophor-El vehicle, while at the same time providing a local depot of high drug concentration for at least four weeks after injection.

**Figure 3.**
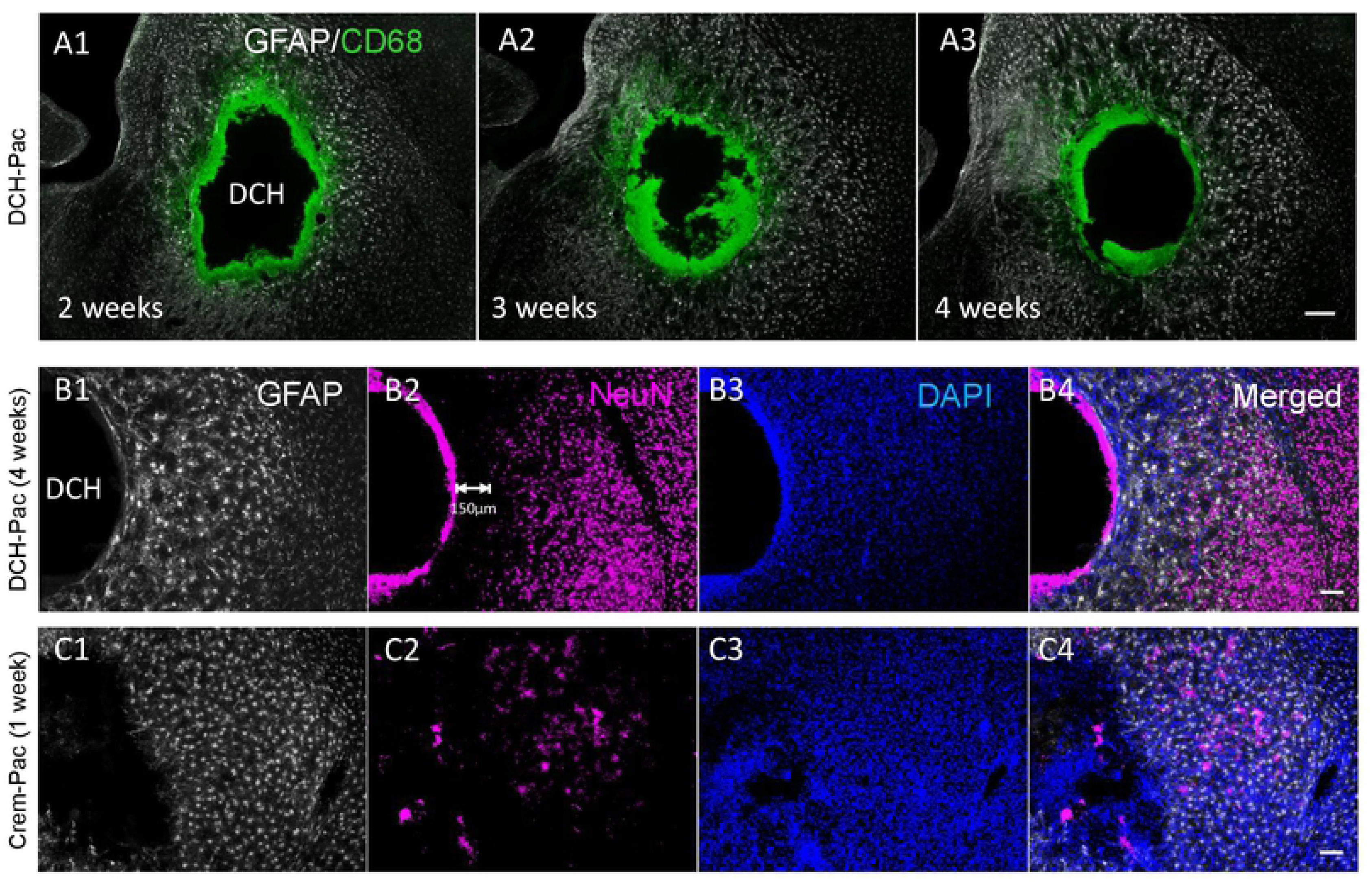
Paclitaxel-DCH depots persist locally for up to 4 weeks *in vivo* and are superior at preserving viable adjacent healthy CNS tissue compared to paclitaxel in Cremophor EL vehicle. **A.** CD68 positive cells are present at the DCH-paclitaxel interface for up to 4 weeks post injection but there is minimal diminution of the depot size, minimal material resorption and minimal infiltration of inflammatory cells into the DCH-paclitaxel depot. **B.** NeuN positive, viable neurons are present in normal density and intermingled with mildly reactive astrocytes in close proximity with the DCH-paclitaxel depot (DCH-Pac). **C.** In contrast, there is pronounced depletion of NeuN positive neurons and severe proliferative reactive astrogliosis in tissue adjacent to injection of paclitaxel in Cremophor EL vehicle (Crem-Pac). Scale bar, 200 µm for all images.

### 3.4. DCH-paclitaxel depots reduce bioluminescence imaging of hGBM cells, delay tumor expansion, and confer a significant survival advantage in NSG mice

Since the DCH-paclitaxel system showed prolonged local availability of active drug in combination with a favorable foreign body response in healthy mice compared to the standard Cremophor EL vehicle, we evaluated its effects on in vivo xenotransplantation model of glioblastoma. We injected a 2 µL suspension of 100,000 cells from a patient-derived gliomasphere line (GFP-positive, Luciferase-positive) into the striatum, followed immediately, but separately, by a second injection of 2 µL of DCH placed directly above the hGBM cells (**Figure 4A).** Animals received either DCH-only or DCH-paclitaxel. We used non-invasive, semi-quantitative bioluminescence imaging to estimate hGBM tumor size and follow the approximate progression of tumor growth in live animals,

**Figure 4:**
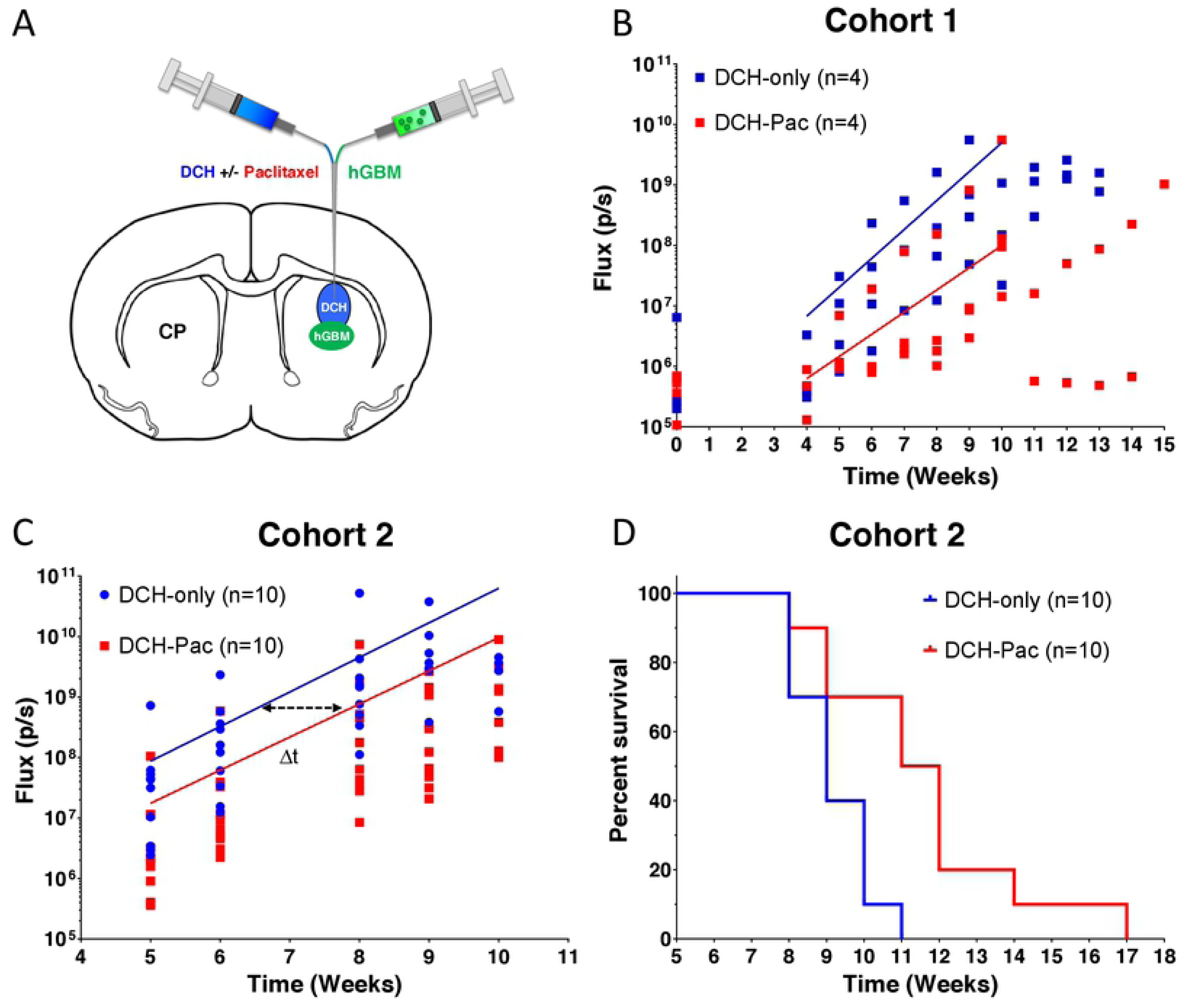
DCH-paclitaxel reduces hGBM growth progression rates monitored by luciferase bioluminescence imaging *in vivo* and prolongs survival. **A.** Schematic of the mouse brain identifying the anatomical location of the gliomasphere and then DCH injections within the caudate putamen (CP) region of the striatum. **B and C**. Graphs showing Luciferase Total Flux (p/s = photons/second) progression as a function of time as measured by bioluminescence IVIS imaging for each animal receiving either DCH only or DCH-paclitaxel in cohort 1 (n=4 animals per group) (**B)** or cohort 2 (n=10 animals per group) **(C)**. An exponential growth regression was applied to each treatment group in **B** and **C** which is represented by a straight line on the semi-log plot. The delta t (Δt) is the time between the same absolute average flux value between the two treatment groups and was calculated to be approximately 1.5 weeks for cohort 2 **D.** Kaplan Meyer survival curve for cohort 2 animals which demonstrated that DHC-paclitaxel conferred a significant median survival time increase of 2.5 week (or a 23% extension of life) compared to the DCH alone (p = 0.0063).

We first conducted a small sample pilot study (Cohort 1, n = 4 animals per group) in which bioluminescence imaging was conducted immediately after brain injections and at weekly intervals from 4 weeks onwards (**Figure 4B**). In both treatment groups, there was no significant increase in the bioluminescence flux signal at 4 weeks compared to several hours after injection. In the hGBM/DCH-only group, there was a measurable increase in the bioluminescence flux signal at 5 weeks, which continued to increase exponentially until 10 weeks when the signal plateaued as the hGBM tumors reached a terminal size and animals required euthanasia (**Figure 4B**). To evaluate the effect of paclitaxel treatment on hGBM cells, we applied exponential regression analysis on the bioluminescence flux signal values across the interval of 5 to 9 weeks of exponential growth for each animal to determine (i) the extrapolated flux signal at the initial time in which tumor growth kinetics appear to begin exponential growth (A_o_); and (ii) the kinetic rate constant of the exponential growth (k). Paclitaxel treatment resulted in (i) an approximately 4-fold reduction in hGBM bioluminescence signal compared to the control group at 4 weeks after injection, which was just prior to the start of measurable increase, and (ii) reduced the subsequent rate of signal increase by approximately 23% (**Figure 4B**).

To test whether the effect of paclitaxel treatment observed in Cohort 1 (**Figure 4B**) was robust, we performed an equivalent study on a second, larger group of animals (Cohort 2, n=10). We conducted bioluminescence imaging over the time period of exponential signal increase from 5 to10 weeks after injection of hGBM cells. In this second, larger group of mice, the DCH-paclitaxel treatment also reduced the initial size of the of hGBM bioluminescence signal by approximately 4-fold, but the rate of the increase in signal was not altered (as shown by the similar gradients for the two curves in **Figure 4C**) and luminescence signals eventually reached similar levels to that of DCH-only controls. As expected, the high variability of luminescence signal among different animals precluded statistical evaluation and the values obtained can only be regarded as semi-quantitative estimates. Nevertheless, the clear shift to the right in the timing of the exponential increase in bioluminescence in the DCH-paclitaxel group (Δ**t in Figure 4C)** suggested that DCH-paclitaxel treatment reduced the number of hGBM cells, but did not eliminate them and for this reason tumor expansion continued at a similar exponential rate but in a delayed manner. This suggestion was further supported by the observation that DCH-paclitaxel treatment was associated with a statistically significant 2.5 longer survival time in DCH-paclitaxel animals compared with DCH-only controls (p value = 0.0063 in log-rank test) (**Figure 4D**), representing a 23% extension of life in this hGBM xenotransplantation model. This prolonged survival time correlated well with the approximately 1.5-week delay in the onset and progression of the exponential increase of bioluminescence imaging (Δt, **Figure 4C**) in DCH-paclitaxel treated mice. Together, these data strongly suggested that our DCH-paclitaxel treatment initially reduced the hGBM tumor load and thereby delayed the expansion of hGBM growth, but eventually the hGBM cells were able to escape treatment.

### DCH-paclitaxel depots reduce local hGBM cell survival but hGBM cells are able to migrate away from treatment zone

Although bioluminescence imaging provided a useful estimate of the effects of DCH-paclitaxel on overall hGBM tumor growth, it could not provide direct information about the effects of DCH-paclitaxel on hGBM cell number or location. To obtain and quantify this type of information, we conducted histological evaluations in particular at 5 weeks after hGBM cell and DCH injection (**Figures 5**), the time point at which hGBM tumors began to exhibit exponential growth as indicated by bioluminescence (**Figure 4B**). hGBM tumor cells were identified by immunofluorescence staining for GFP and human nuclear antigen (NA) antibodies.

**Figure 5.**
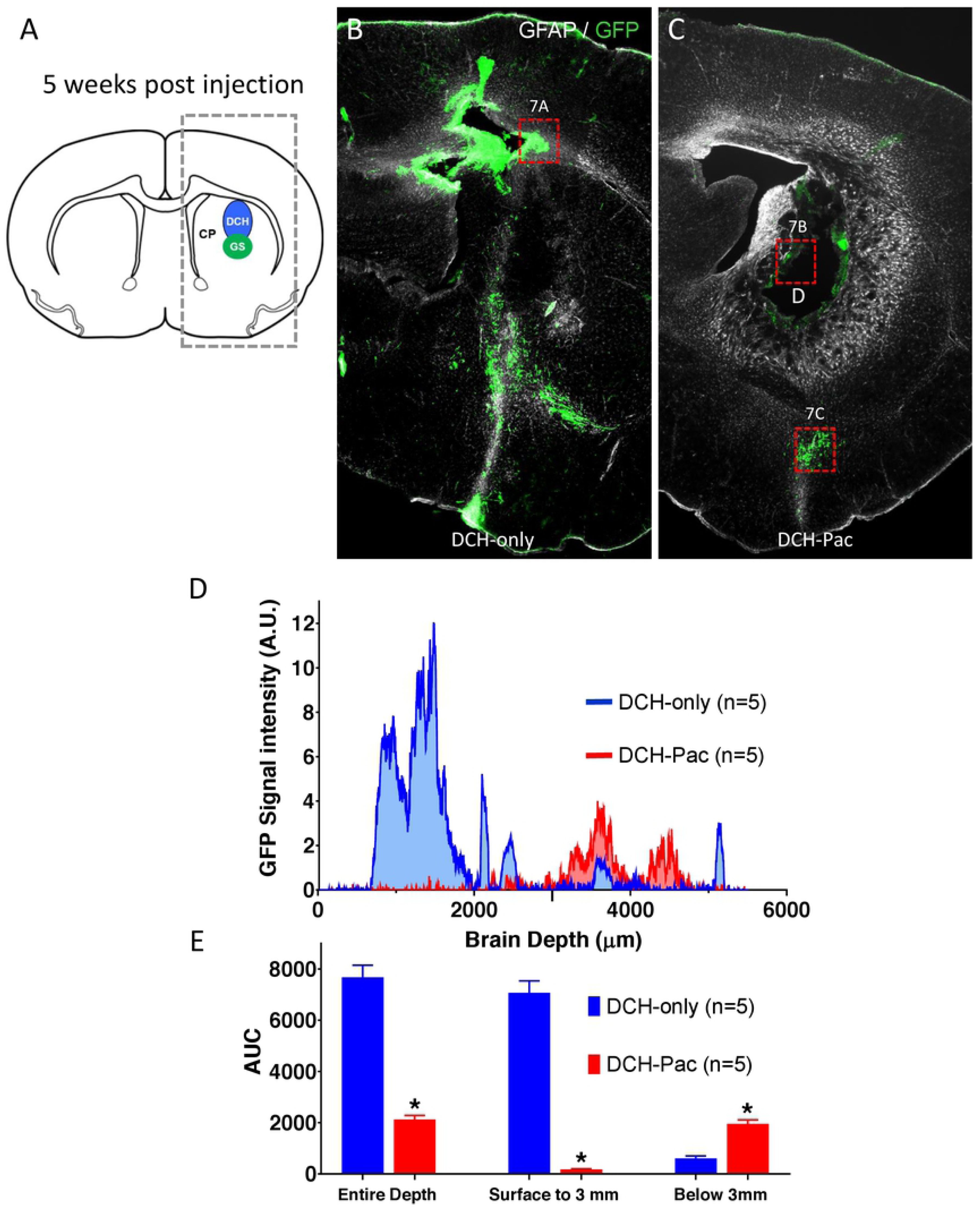
The DCH-paclitaxel system substantively ablates locally residing hGBM cells but does not prevent the migration of tumor cells within brain parenchyma. **A.** Schematic of mouse forebrain showing the location of hGBM and DCH injections in the caudate putamen (CP). Box of dashed lines delineates the area quantitatively analyzed for the presence of GFP labeled cells in 5-week post injection tissue and presented in D and E. **B,C.** Survey immunofluorescent images of staining for GFP (hGBM cells) and GFAP show the location of hGBM cells in forebrain at 5 weeks post injection. **B.** Mouse that received hGBM cells and DCH-only. Note the high density of GFP-positive hGBM cells immediately above and below the injection site. **C.** Mouse that received hGBM cells and DCH-paclitaxel. Note the essential absence of GFP-positive hGBM cells immediately around the persisting DCH-paclitaxel depot (D). Note also the presence of hGBM cells that have migrated away from the injection site (box shown at higher magnification in 7C). **D.** Graph of quantification of GFP signal from hGBM cells. GFP staining intensity was measured as a function of brain depth in serial linear units in the boxed area shown in **A**. As expected, animals receiving DCH-only (blue) exhibited a high intensity of GFP signal in and immediately above the injection region (0.5 to 3 mm). In contrast, animals receiving DCH-paclitaxel (DCH-Pac, red) exhibited little or no GFP signal in this area, but did exhibit substantive signal at deeper levels (around 4mm). **E**. Bar graphs quantifying area under the Curve (AUC) for GFP staining intensity. Over the entire depth of the brain, DCH-paclitaxel treated animals (DCH-Pac, red) exhibited an over 72% reduction in hGBM-derived GFP signal compared with DCH-only (p<0.001). In the area immediately around the hGBM and DCH injections (surface to 3 mm), DCH-paclitaxel treated animals exhibited an over 97% reduction in hGBM-derived GFP signal (p<0.001). In contrast, in the deeper areas away from DCH depots (around 4 mm), DCH-paclitaxel treated animals exhibited a 65% greater hGBM-derived GFP signal (p<0.001). n = 5 per group for all measures.

Control animals receiving DCH-only consistently exhibited a focal large cluster of GFP-positive and HNA-positive cells in the dorsal striatum and cerebral cortex along the needle track above the striatal injection site, which represented the predominant hGBM cell population (**Figure 5A,B**). In addition, many hGBM cells had migrated along disrupted tissue planes and white matter tracts into deep subcortical regions and the contralateral hemisphere (**Figure 5B**). By contrast, DCH-paclitaxel treated animals showed only occasional hGBM cells in the dorsal striatum and overlying cerebral cortex while the bulk of GFP-positive and HNA signal in these treatment animals was observed in two main regions (i) at the margins of the DCH deposit and (ii) in deep subcortical regions well below the DCH deposit site (**Figure 5C**).

To quantify the number of hGBM cells in different brain locations at 5 weeks after injection we measured GFP staining intensity as a function of brain depth in serial linear units relative to the cortical surface at the injection coordinates (**Figure 5D**). These measurements showed that the DCH-paclitaxel treatment significantly reduced the number of hGBM cells in the area immediately around the injection site (i.e. cortical surface to 3 mm deep) by 97% (p<0.001) (**Figure 5D,E**). The number of total tumor cells across the entire depth of the brain was significantly reduced by approximately 72.3% (p<0.001) in the paclitaxel group compared to controls (**Figure 5D,E**). Interestingly, in ventral brain regions 1 or more mm distal from the injection site (**Figure 5C**), the number of tumor cells was somewhat higher in the paclitaxel group (**Figure 5D,E**), suggesting that tumor cells had migrated away from the paclitaxel.

## DISCUSSION

Extent of resection (EOR) has been highly correlated to improved survival in low and high-grade glioma [29–32]. However, the ability to perform near complete tumor resection is dependent on the location of the primary mass within the brain. Tumors located in deep sub-cortical or midline structures, in close proximity to ventricles, near large vessels or at eloquent brain regions, are often classified as inoperable or unsuitable for large volume tumor resection due to the high risk of severe postsurgical morbidity[33, 34]. These unresectable tumors are associated with shorter patient survival[32, 33]. The paclitaxel-loaded DCH system represents a potential approach for debulking these difficult to access tumors without causing significant post-surgical morbidity.

In this study we show that DCH can locally deliver a potent hydrophobic chemotherapeutic agent, paclitaxel, to a focal site within the brain and substantively ablate locally residing hGBM tumor cells while minimizing the extent of damage to healthy host tissue. Although DCH-paclitaxel treatment massively depleted the number of local hGBM cells, the surviving hGBM cells were able to avoid the treatment area and migrate to deeper areas of the brain. This finding highlights the importance of using realistic cellular models. Previous studies using serum-derived glioma cells or utilizing in vitro or subcutaneous environments under-estimate the effect of cell migration and thus likely overstate their efficacy. Despite the difficulty of targeting migrating tumor cells with a focal treatment vehicle, the potential survival benefit of this technology should warrant further investigation and consideration.

One potential solution to the problem of cell migration involves a combination of cell homing and cell ablation in which chemoattractants could be used to guide migration of glioblastoma cells towards a focal “kill zone” where they are destroyed by chemotherapeutic agents [35]. Many chemoattractant molecules and other physical cell guidance systems are being identified, including various canonical cytokines and growth factors [36]. Another potential use of this technology involves pairing it with other minimally invasive tumor ablation techniques. Devices using ultrasound [37] or thermo-ablation [38] are being developed to debulk the central tumor mass. These technologies leave a hole in the tumor mass that could potentially be filled with a hydrogel for extended tumor control.

One problem with delivering chemotherapeutic agents at the time of surgery is the concern for wound healing. The initial trials with gliadel were associated with an increase in wound break-down. It is for this reason, that many systemic chemotherapies are started several weeks after the surgery to allow for the wound to heal. One advantage of minimally invasive injection techniques is that they involve smaller wounds, which are less prone to wound complications and break-down.

In conclusion, this study demonstrates the potential for using DCH to deliver a potent chemotherapeutic agent to a focal site within the brain and effectively chemically debulk a central area of tumor with a resulting survival benefit. This technology brings hope to a population of patients for whom surgical debulking is not possible.

## Author Contributions

T.M.O’S. and M.C.G. contributed equally to this work;

T.M.O’S., M.C.G., T.J.D., H.I.K., M.V.S. designed experiments;

T.M.O’S., M.C.G., A.M.B., A.L.W., D.H., H.S. conducted experiments;

T.M.O’S, M.C.G., T.J.D., H.I.K., M.V.S. analyzed data;

T.M.O’S, M.C.G.,T.J.D., H.I.K., M.V.S. N.M., B.S. prepared the manuscript.

## ACKNOWLEDGEMENTS

We thank the Microscopy Core Resource of the UCLA Broad Stem Cell Research Center-CIRM Laboratory. This work was supported by the Dr. Miriam and Sheldon G. Adelson Medical Foundation (M.V.S., T.J.D. and H.I.K.), the US National Institutes of Health (NS084030 to M.V.S. and NS053563 to H.I.K), and a training grant from the California Institute for Regenerative Medicine (M.C.G.).

## COMPLIANCE WITH ETHICAL STANDARDS

Author MCG declares he has no conflicts of interest

Author ALW declares he has no conflicts of interest

Author AMC declares he has no conflicts of interest

Author TMO declares he has no conflicts of interest

Author DH declares he has no conflicts of interest

Author NM declares he has no conflicts of interest

Author BS declares she has no conflicts of interest

Author HS declares he has no conflicts of interest

Author TJ declares he has no conflicts of interest

Author MVS declares he has no conflicts of interest

Author HIK declares he has no conflicts of interest

## REFERENCES

1. Wen PY, Kesari S (2008) Malignant gliomas in adults. The New England journal of medicine 359: 492–507 doi:10.1056/NEJMra0708126

2. Chaichana KL, Zadnik P, Weingart JD, Olivi A, Gallia GL, Blakeley J, Lim M, Brem H, Quinones-Hinojosa A (2013) Multiple resections for patients with glioblastoma: prolonging survival. Journal of neurosurgery 118: 812–820 doi:10.3171/2012.9.JNS1277

3. Gruber ML, Hochberg FH (1990) Systematic evaluation of primary brain tumors. Journal of nuclear medicine: official publication, Society of Nuclear Medicine 31: 969–971

4. Sanai N, Berger MS (2011) Extent of resection influences outcomes for patients with gliomas. Revue neurologique 167: 648–654 doi:10.1016/j.neurol.2011.07.004

5. Yonezawa H, Hirano H, Uchida H, Habu M, Hanaya R, Oyoshi T, Sadamura Y, Hanada T, Tokimura H, Moinuddin F, Arita K (2017) Efficacy of bevacizumab therapy for unresectable malignant glioma: A retrospective analysis. Molecular and clinical oncology 6: 105–110 doi:10.3892/mco.2016.1086

6. Stewart LA (2002) Chemotherapy in adult high-grade glioma: a systematic review and meta-analysis of individual patient data from 12 randomised trials. Lancet 359: 1011–1018

7. Stupp R, Mason WP, van den Bent MJ, Weller M, Fisher B, Taphoorn MJ, Belanger K, Brandes AA, Marosi C, Bogdahn U, Curschmann J, Janzer RC, Ludwin SK, Gorlia T, Allgeier A, Lacombe D, Cairncross JG, Eisenhauer E, Mirimanoff RO, European Organisation for R, Treatment of Cancer Brain T, Radiotherapy G, National Cancer Institute of Canada Clinical Trials G (2005) Radiotherapy plus concomitant and adjuvant temozolomide for glioblastoma. The New England journal of medicine 352: 987–996 doi:10.1056/NEJMoa043330

8. Johnson BE, Mazor T, Hong C, Barnes M, Aihara K, McLean CY, Fouse SD, Yamamoto S, Ueda H, Tatsuno K, Asthana S, Jalbert LE, Nelson SJ, Bollen AW, Gustafson WC, Charron E, Weiss WA, Smirnov IV, Song JS, Olshen AB, Cha S, Zhao Y, Moore RA, Mungall AJ, Jones SJM, Hirst M, Marra MA, Saito N, Aburatani H, Mukasa A, Berger MS, Chang SM, Taylor BS, Costello JF (2014) Mutational analysis reveals the origin and therapy-driven evolution of recurrent glioma. Science 343: 189–193 doi:10.1126/science.1239947

9. Kitange GJ, Carlson BL, Schroeder MA, Grogan PT, Lamont JD, Decker PA, Wu W, James CD, Sarkaria JN (2009) Induction of MGMT expression is associated with temozolomide resistance in glioblastoma xenografts. Neuro-oncology 11: 281–291 doi:10.1215/15228517-2008-090

10. Chang SM, Kuhn JG, Robins HI, Schold SC, Jr., Spence AM, Berger MS, Mehta M, Pollack IF, Rankin C, Prados MD (2001) A Phase II study of paclitaxel in patients with recurrent malignant glioma using different doses depending upon the concomitant use of anticonvulsants: a North American Brain Tumor Consortium report. Cancer 91: 417–422

11. Moses MA, Brem H, Langer R (2003) Advancing the field of drug delivery: taking aim at cancer. Cancer cell 4: 337–341

12. Rowland MJ, Parkins CC, McAbee JH, Kolb AK, Hein R, Loh XJ, Watts C, Scherman OA (2018) An adherent tissue-inspired hydrogel delivery vehicle utilised in primary human glioma models. Biomaterials 179: 199–208 doi:10.1016/j.biomaterials.2018.05.054

13. Puente P, Fettig N, Luderer MJ, Jin A, Shah S, Muz B, Kapoor V, Goddu SM, Salama NN, Tsien C, Thotala D, Shoghi K, Rogers B, Azab AK (2018) Injectable Hydrogels for Localized Chemotherapy and Radiotherapy in Brain Tumors. Journal of pharmaceutical sciences 107: 922–933 doi:10.1016/j.xphs.2017.10.042

14. Cui Y, Xu Q, Chow PK, Wang D, Wang CH (2013) Transferrin-conjugated magnetic silica PLGA nanoparticles loaded with doxorubicin and paclitaxel for brain glioma treatment. Biomaterials 34: 8511–8520 doi:10.1016/j.biomaterials.2013.07.075

15. Zhao M, Danhier F, Bastiancich C, Joudiou N, Ganipineni LP, Tsakiris N, Gallez B, Rieux AD, Jankovski A, Bianco J, Preat V (2018) Post-resection treatment of glioblastoma with an injectable nanomedicine-loaded photopolymerizable hydrogel induces long-term survival. International journal of pharmaceutics 548: 522–529 doi:10.1016/j.ijpharm.2018.07.033

16. Tyler B, Fowers KD, Li KW, Recinos VR, Caplan JM, Hdeib A, Grossman R, Basaldella L, Bekelis K, Pradilla G, Legnani F, Brem H (2010) A thermal gel depot for local delivery of paclitaxel to treat experimental brain tumors in rats. Journal of neurosurgery 113: 210–217 doi:10.3171/2009.11.JNS08162

17. Hemmati HD, Nakano I, Lazareff JA, Masterman-Smith M, Geschwind DH, Bronner-Fraser M, Kornblum HI (2003) Cancerous stem cells can arise from pediatric brain tumors. Proceedings of the National Academy of Sciences of the United States of America 100: 15178–15183 doi:10.1073/pnas.2036535100

18. Laks DR, Crisman TJ, Shih MY, Mottahedeh J, Gao F, Sperry J, Garrett MC, Yong WH, Cloughesy TF, Liau LM, Lai A, Coppola G, Kornblum HI (2016) Large-scale assessment of the gliomasphere model system. Neurooncology 18: 1367–1378 doi:10.1093/neuonc/now045

19. Dull T, Zufferey R, Kelly M, Mandel RJ, Nguyen M, Trono D, Naldini L (1998) A third-generation lentivirus vector with a conditional packaging system. Journal of virology 72: 8463–8471

20. Zhang S, Anderson MA, Ao Y, Khakh BS, Fan J, Deming TJ, Sofroniew MV (2014) Tunable diblock copolypeptide hydrogel depots for local delivery of hydrophobic molecules in healthy and injured central nervous system. Biomaterials 35: 1989–2000 doi:10.1016/j.biomaterials.2013.11.005

21. Lee SH, Yoo SD, Lee KH (1999) Rapid and sensitive determination of paclitaxel in mouse plasma by high-performance liquid chromatography. Journal of chromatography B, Biomedical sciences and applications 724: 357–363

22. Faulkner JR, Herrmann JE, Woo MJ, Tansey KE, Doan NB, Sofroniew MV (2004) Reactive astrocytes protect tissue and preserve function after spinal cord injury. The Journal of neuroscience: the official journal of the Society for Neuroscience 24: 2143–2155 doi:10.1523/JNEUROSCI.3547-03.2004

23. Lee SY (2016) Temozolomide resistance in glioblastoma multiforme. Genes & diseases 3: 198–210 doi:10.1016/j.gendis.2016.04.007

24. Hafner M, Niepel M, Chung M, Sorger PK (2016) Growth rate inhibition metrics correct for confounders in measuring sensitivity to cancer drugs. Nat Meth 13: 521–527 doi:10.1038/nmeth.3853 http://www.nature.com/nmeth/journal/v13/n6/abs/nmeth.3853.html - supplementary-information

25. Yang C-Y, Song B, Ao Y, Nowak AP, Abelowitz RB, Korsak RA, Havton LA, Deming TJ, Sofroniew MV (2009) Biocompatibility of amphiphilic diblock copolypeptide hydrogels in the central nervous system. Biomaterials 30: 2881–2898 doi:10.1016/j.biomaterials.2009.01.056

26. Rath BH, Fair JM, Jamal M, Camphausen K, Tofilon PJ (2013) Astrocytes Enhance the Invasion Potential of Glioblastoma Stem-Like Cells. PLoS ONE 8: e54752 doi:10.1371/journal.pone.0054752

27. Sin WC, Aftab Q, Bechberger JF, Leung JH, Chen H, Naus CC (2016) Astrocytes promote glioma invasion via the gap junction protein connexin43. Oncogene 35: 1504–1516 doi:10.1038/onc.2015.210

28. Placone AL, Quinones-Hinojosa A, Searson PC (2016) The role of astrocytes in the progression of brain cancer: complicating the picture of the tumor microenvironment. Tumour biology: the journal of the International Society for Oncodevelopmental Biology and Medicine 37: 61–69 doi:10.1007/s13277-015-4242-0

29. Michel Lacroix, Dima Abi-Said, Daryl R. Fourney, Ziya L. Gokaslan, Weiming Shi, Franco DeMonte, Frederick F. Lang, Ian E. McCutcheon, Samuel J. Hassenbusch, Eric Holland, Kenneth Hess, Christopher Michael, Daniel Miller, Raymond Sawaya (2001) A multivariate analysis of 416 patients with glioblastoma multiforme: prognosis, extent of resection, and survival. Journal of Neurosurgery 95: 190–198 doi:doi:10.3171/jns.2001.95.2.0190

30. Stummer W, Reulen H-J, Meinel T, Pichlmeier U, Schumacher W, Tonn J-C, Rohde V, Oppel F, Turowski B, Woiciechowsky C, Franz K, Pietsch T, Group A-GS (2008) EXTENT OF RESECTION AND SURVIVAL IN GLIOBLASTOMA MULTIFORME: IDENTIFICATION OF AND ADJUSTMENT FOR BIAS. Neurosurgery 62: 564–576 doi:10.1227/01.neu.0000317304.31579.17

31. Nader Sanai, Mei-Yin Polley, Michael W. McDermott, Andrew T. Parsa, Mitchel S. Berger (2011) An extent of resection threshold for newly diagnosed glioblastomas. Journal of Neurosurgery 115: 3–8 doi:doi:10.3171/2011.2.JNS10998

32. Kuhnt D, Becker A, Ganslandt O, Bauer M, Buchfelder M, Nimsky C (2011) Correlation of the extent of tumor volume resection and patient survival in surgery of glioblastoma multiforme with high-field intraoperative MRI guidance. Neuro-Oncology 13: 1339–1348 doi:10.1093/neuonc/nor133

33. Orringer D, Lau D, Khatri S, Zamora-Berridi GJ, Zhang K, Wu C, Chaudhary N, Sagher O (2012) Extent of resection in patients with glioblastoma: limiting factors, perception of resectability, and effect on survival. J Neurosurg 117: 851–859 doi:10.3171/2012.8.jns12234

34. Kubben PL, ter Meulen KJ, Schijns OE, ter Laak-Poort MP, van Overbeeke JJ, van Santbrink H (2011) Intraoperative MRI-guided resection of glioblastoma multiforme: a systematic review. The Lancet Oncology 12: 1062–1070 doi:10.1016/s1470-2045(11)70130-9

35. van der Sanden B, Appaix F, Berger F, Selek L, Issartel J-P, Wion D (2013) Translation of the ecological trap concept to glioma therapy: the cancer cell trap concept. Future Oncology 9: 817–824 doi:10.2217/fon.13.30

36. Kim Y (2013) Regulation of cell proliferation and migration in glioblastoma: new therapeutic approach. Frontiers in oncology 3: 53 doi:10.3389/fonc.2013.00053

37. Hersh DS, Kim AJ, Winkles JA, Eisenberg HM, Woodworth GF, Frenkel V (2016) Emerging Applications of Therapeutic Ultrasound in Neuro-oncology: Moving Beyond Tumor Ablation. Neurosurgery 79: 643–654 doi:10.1227/NEU.0000000000001399

38. Patel P, Patel NV, Danish SF (2016) Intracranial MR-guided laser-induced thermal therapy: singlecenter experience with the Visualase thermal therapy system. Journal of neurosurgery 125: 853–860 doi:10.3171/2015.7.JNS15244

